# Trends and spatial distribution of animal bites and vaccination status among victims and the animal population, Uganda: A veterinary surveillance system analysis, 2013 – 2017

**DOI:** 10.1101/852517

**Authors:** Fred Monje, Daniel Kadobera, Deo Birungi Ndumu, Lilian Bulage, Alex Riolexus Ario

## Abstract

**Background:** Rabies is a vaccine-preventable fatal zoonotic disease. Uganda, through the veterinary surveillance system at National Animal Disease Diagnostics and Epidemiology Centre (NADDEC), captures animal bites (a proxy for rabies) on a monthly basis from districts. We established trends of incidence of animal bites and corresponding post-exposure prophylactic anti-rabies vaccination in humans (PEP), associated mortality rates in humans, spatial distribution of animal bites, and pets vaccinated during 2013-2017.

**Methods:** We reviewed rabies surveillance data at NADDEC from 2013-2017. The surveillance system captures persons reporting bites by a suspected rabid dog/cat/wild animal, human deaths due to suspected rabies, humans vaccinated against rabies, and pets vaccinated. Number of total pets was obtained from the Uganda Bureau of Statistics. We computed incidence of animal bites and corresponding PEP in humans, and analyzed overall trends, 2013-2017. We also examined human mortality rates and spatial distribution of animal bites/rabies and pets vaccinated against rabies.

**Findings:** We identified 8,240 persons reporting animal bites in Uganda during 2013-2017; overall incidence of 21 bites/100,000population. The incidence significantly decreased from 8.2/100,000 in 2013 to 1.1/100,000 in 2017 (OR=0.62, p= 0.0074). Of the 8,240 persons with animal bites, 6,799 (82.5%) received PEP, decreasing from 94% in 2013 to 71% in 2017 (OR=0.65, p<0.001). Among 1441 victims, who reportedly never received PEP, 156 (11%) died. Western region had a higher incidence of animal bites (37/100,000) compared to other regions. Only 5.6% (124,555/2,240,000) of all pets in Uganda were vaccinated. There was a decline in the reporting rate of animal bites.

**Conclusion:** While reported animal bites decreased in Uganda, so did PEP among humans. Very few pets received anti-rabies vaccine. Evaluation of barriers to complete reporting may facilitate interventions to enhance surveillance quality. We recommended improved vaccination of pets against rabies, and immediate administration of exposed humans with PEP.

**Author Summary:** Rabies is a deadly viral disease, that is transmitted mainly by dog bites. Globally at least 59,000 deaths are reported to occur annually- mostly in Sub-Saharan Africa and Asia. However, rabies can be prevented through vaccination of pets (dogs and cats) and administration of rabies vaccine in humans exposed to rabies. In our study we reviewed secondary data of animal bites and rabies captured at the National Animal disease diagnostic epidemiology centre in Entebbe for the period 2013-2017. We found that of 1441 animal bite victims who never received rabies vaccine, only 156 (11%) died hence need for immediate administration of exposed humans with rabies and sensitization of the public about the consequences of animal bites and need for urgent health care. There was a decline in the reporting rate of animal bites during the study period suggesting that evaluation of the barriers to complete reporting may facilitate interventions to enhance surveillance quality. Less than 10% of the pets in the Uganda were vaccinated against rabies hence need for improved vaccination of pets against rabies through appropriated legislation.

## Introduction

Rabies is a fatal viral vaccine-preventable zoonotic disease that can infect all mammals (1–3). Worldwide, canine rabies causes at least 59,000 human deaths and 8.6 billion USD (95% CIs: 2.9-21.5 billion) in economic losses annually (3). Rabies is transmitted primarily through bites from an infected rabid animal; however, it can also be transmitted from licks or scratches from an infected rabid animal or, rarely, through transplantation of tissues or organs from an infected individual (4–6). Rabies occurs on all continents except Antarctica, with most cases reported in Africa and Asia (7,8). Almost all cases of human rabies are due to bites from infected dogs (7). In dogs, the incubation period of rabies is 10 days to 6 months, with most cases manifesting signs between 2 weeks to 3 months after exposure (9). In humans, the average incubation period of rabies is 20-60 days, though it can last up to several years (6,10). Following exposure, suggestive signs of rabies in humans include intense pruritus, beginning at the site of the bite and progressing to involve the limb or side of the face, myoedema, neurological symptoms including confusion and aggression, partial paralysis, involuntary muscle twitching, rigid neck muscles, convulsions, hyperventilation and difficulty breathing, hypersalivation, hydrophobia, difficulty swallowing, hallucinations, nightmares, and photophobia. As disease progresses, the patient enters a coma and dies (6).

Prevention of rabies in animals is primarily achieved through vaccination. Indeed, in developed countries, mass canine vaccination coupled with oral vaccination in wildlife have greatly contributed to the elimination of rabies in canines, and consequently a reduction in human rabies (7). In humans, rabies prevention typically occurs through rabies Post-Exposure Prophylaxis (rPEP) in the form of a rabies vaccine. This vaccine should be administered to victims of bites from suspected rabid animals as soon as possible, and continued while the animal is being observed for 10-14 days or pending the results of laboratory tests (6). However, rPEP requires multiple doses, is not always available, and must be administered in a timely manner to be effective. The most cost-effective method of controlling rabies is to prioritize canine vaccination, rather than using reactive rPEP in humans (7).

In Uganda, Ministry of Agriculture, Animal Industry and Fisheries (MAAIF) procures anti-rabies vaccine annually to control rabies in animals, though in limited doses (11). This vaccine is provided to districts based on the magnitude of reported animal bites, the dog population in the district, and confirmed rabies cases in pets. Historically, MAAIF has allocated about 2,000 doses of anti-rabies vaccine for pets to each of the districts in Uganda affected by rabies. This is generally insufficient for the estimated pet populations in each of the districts. While people are able to obtain vaccine from private providers, the cost ($8-12 USD per dose) is prohibitive for most (12).

Among humans, an animal bite is treated as being from a rabid animal, unless the animal can be confirmed to not have rabies (or is incapable of carrying rabies). Surveillance data of animal bites among humans provides vital information to guide resource allocation in the control and prevention of rabies (13). An analysis of human rabies surveillance data from the Epidemiology and Surveillance Division (ESD) of the Ministry of Health in 2001-2015 in Uganda revealed 208,720 animal bites, with 486 suspected human rabies deaths (14). However, Uganda also captures rabies-related data in a veterinary surveillance system, at National Animal Disease Diagnostic Epidemiology Centre (NADDEC). NADDEC is mandated to participate in animal disease monitoring and surveillance, including rabies. Beyond capturing animal bites and suspected human rabies deaths, NADDEC also captures the number of pets vaccinated against rabies (including by private providers) and conducts confirmatory tests of suspected rabid animals. We used data from NADDEC to describe trends in the incidence of animal bites and corresponding rPEP in humans, mortality associated with animal bites in humans, spatial distribution of animal bites, and pets vaccinated in Uganda from 2013-2017.

## Methods

### Study setting and design

We conducted the study using national surveillance data from National Animal Disease Diagnostics and Epidemiology Centre (NADDEC). When a bite from a suspected rabid animal is reported in the communities, Uganda guidelines state that it must be immediately reported by phone to the nearest veterinary officer or animal husbandry officer at the sub-county headquarters. The officer visits the scene of the incident to ensure that the animal (usually dog/cat) is killed and its head packaged for shipment to NADDEC for analysis. Meanwhile, the veterinary officer/animal husbandry officer assesses the circumstances under which the victim was bitten and the vaccination history of the dog/cat at the time of the incident, and writes a referral letter for the victim to receive rPEP at the nearest health facility. A monthly sub-county report with all the animal bites is compiled by the officer and shared with the District Veterinary Officer (DVO), who then compiles all animal bites in the district and enters the data into a standard veterinary surveillance form. The DVO then submits a district monthly veterinary surveillance report to the Commissioner for Animal Health (CAH), which is then forwarded to NADDEC for compilation. Rabies-specific variables captured at NADDEC include date, district name, animal type, animal species, number of humans bitten by suspected rabid animals, survival status of the bitten human, rPEP status of the bitten human, whether or not the pets (dogs/cats) were vaccinated, and whether or not they were destroyed. The findings from veterinary disease data are intended to be shared with the districts through annual DVO meetings (15).

We carried out a retrospective descriptive study involving review of rabies data captured at NADDEC during 2013-2017. We excluded all records that had had missing key variables. To understand more about the completeness of reporting, we also captured information about the number of monthly surveillance reports expected in a year at NADDEC, and number of reports actually received.

We reviewed NADDEC laboratory records for rabies testing for suspected animals. Rabies diagnosis was conducted on animal brain tissue using the direct fluorescent antibody test (dFAT, Onderstepoort Veterinary Institute, Pretoria, South Africa) at NADDEC (16).

### Data management and analysis

We extracted monthly epidemiological rabies data from NADDEC into Microsoft Excel for analysis. Regional and national human population data were obtained from the Uganda National Census 2014. We calculated annual and overall incidence of animal bites by using number of new cases as a numerator and total human population at risk as a denominator during 2013-2017 We drew maps using Quantum Geographic information system (QGIS) to demonstrate geographical distribution of animal bites by district. From the NADDEC data, we computed the annual trends in rPEP and mortality rates in humans associated with animal bites in the period 2013 to 2017. We also computed the proportion of pets vaccinated against rabies during the period 2013-2017, using pet population data obtained from the Uganda Bureau of Statistics from a 2008 census (17)

## Results

We identified 8,240 reports of animal bites in Uganda during the study period, with an overall incidence of 21 animal bites per 100,000 population. The annual incidence significantly decreased from 8.2/100,000 in 2013 to 1.1/100,000 in 2017 (OR=0.62, p=0.0074) (Fig 1). Animal bites were distributed throughout Uganda (Fig 2). Western region had the highest incidence of animal bites (37/100,000), followed by Northern region (22/100,000), Central region (17/100,000), and Eastern region (17/100,000). These differences were largely driven by differences in reported bites during 2013 and 2014; reported incidences were much lower among all regions during 2015-2017 (Fig 3). Laboratory data indicated that 36% (28/77) of the brain samples from suspected rabid animals tested positive for rabies during the study period.

**Fig 1.**
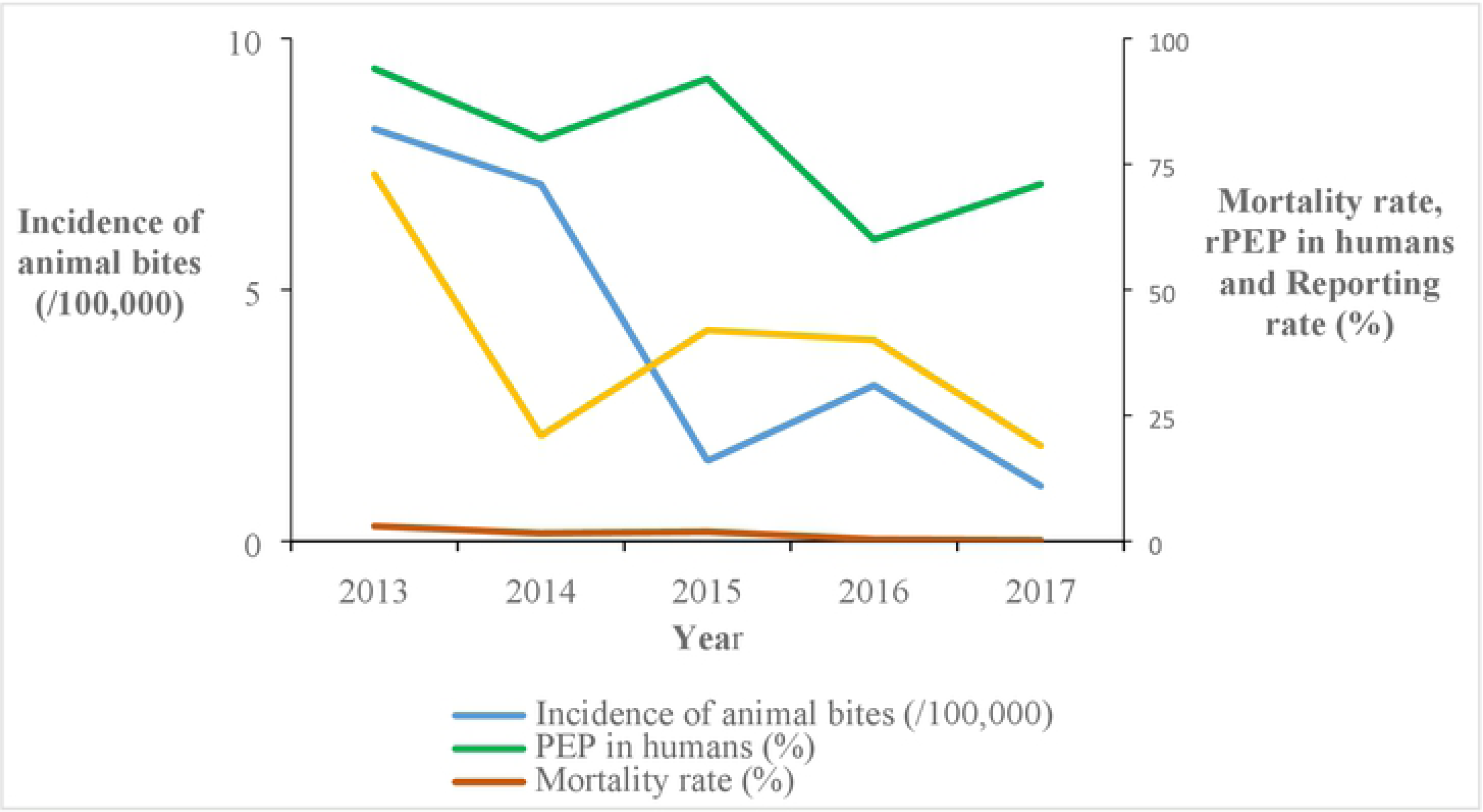
Trends of incidence of animal bites, anti-rabies vaccinations in humans, mortality rates and reporting rates in Uganda: 2013-2017.

**Fig 2.**
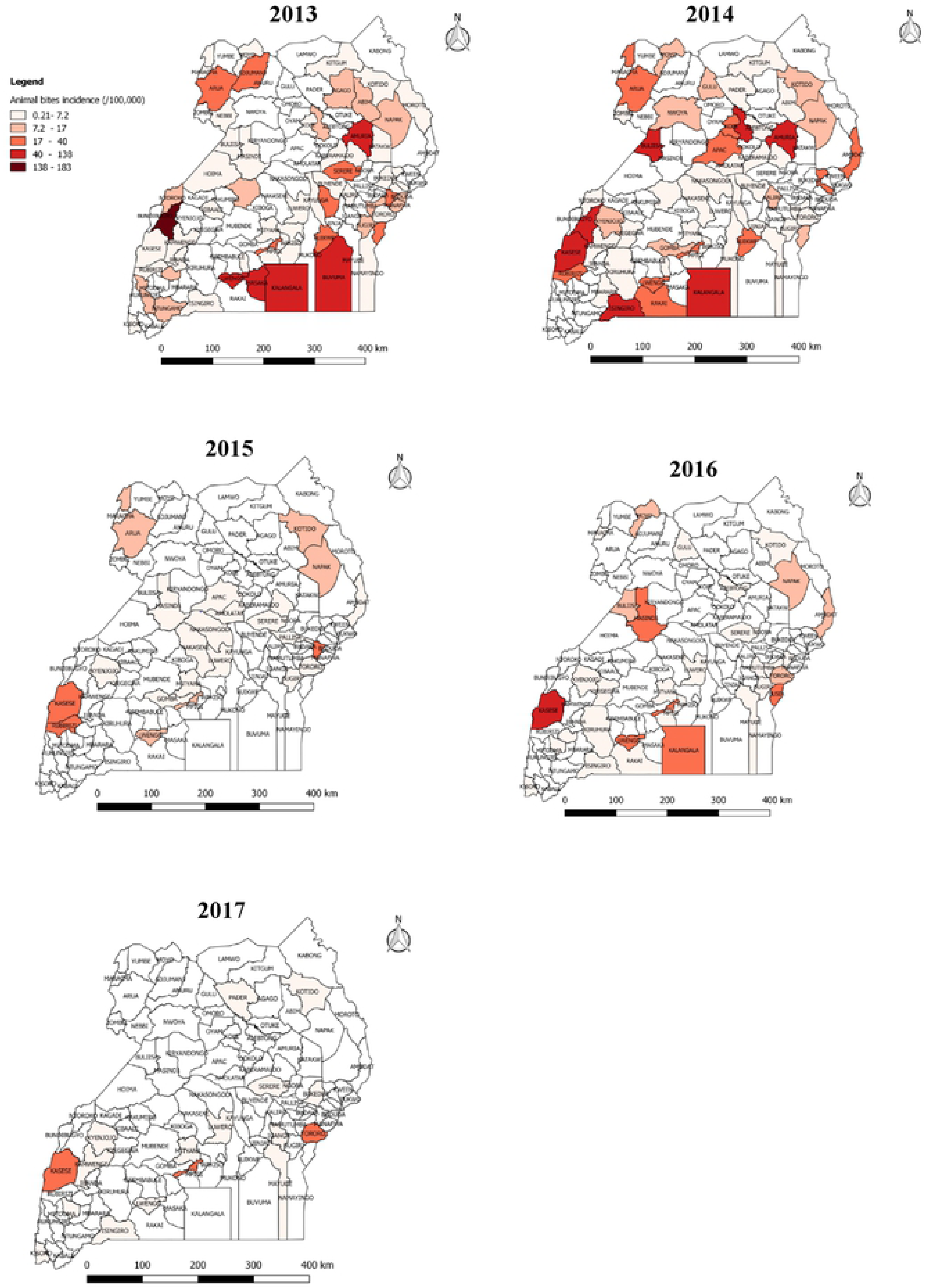
Reported incidence (/100,000 population) of animal bites in Uganda, 2013-2017; Source: Created using QGIS version 2.18.19 from animal bites data, 2013-2017 used in this study.

**Fig 3.**
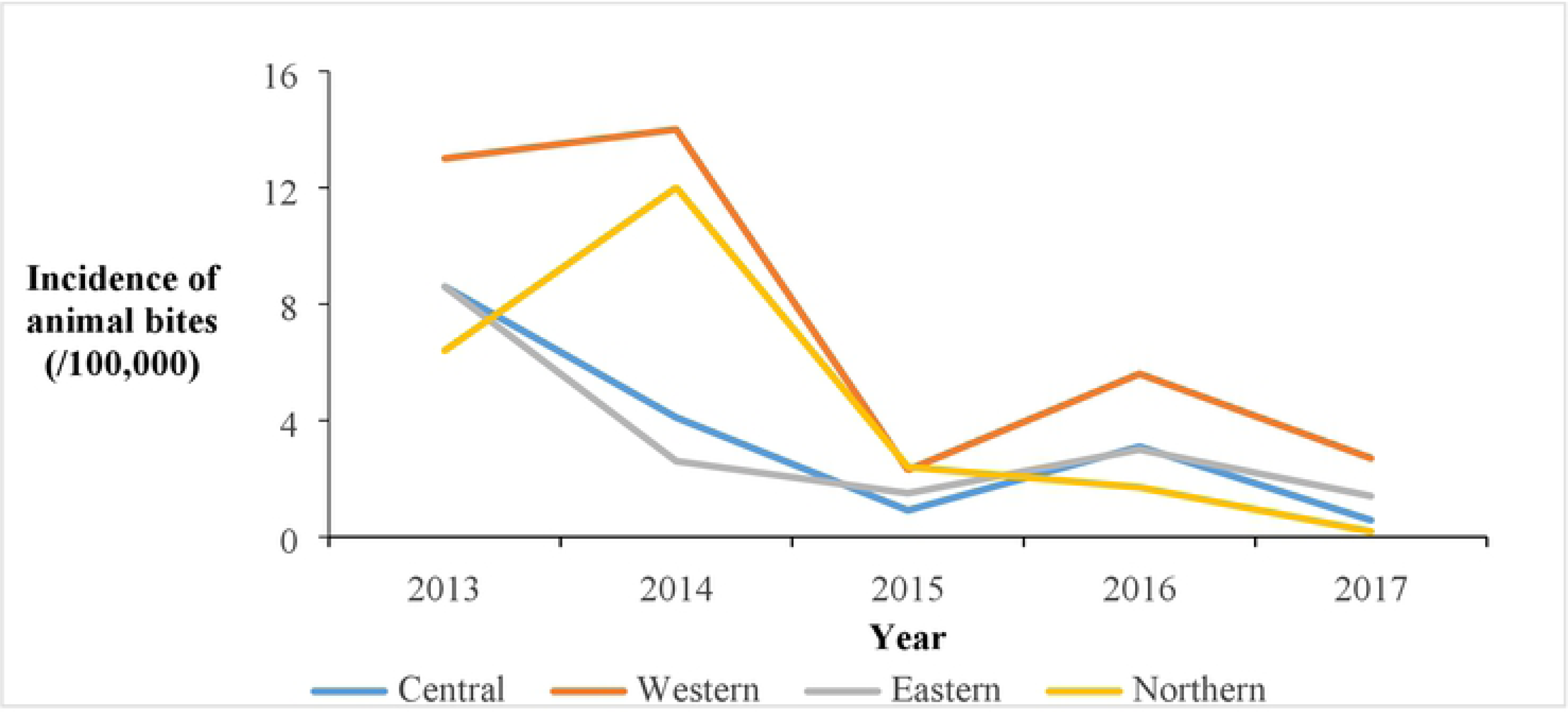
Reported incidence of animal bites (/100,000) by region in Uganda: 2013-2017.

Of the 8,240 human animal bite victims, 6,799 (82.5%) reportedly obtained rPEP. The percentage obtaining rPEP after the bite decreased from 94% in 2013 to 71% in 2017 (OR=0.65; p<0.001) (Figure 1). Among all human animal bite victims, 156 (1.9%) died. The annual reported mortality rates among bitten humans decreased from 3.0% in 2013 to 0.21% in 2017 (OR=0.58, p=0.08) (Figure 1). Of the 2,240,000 pets, 6,997 (0.31%) were reported destroyed by the communities following infliction of bite injuries to humans; the proportion reported destroyed by the communities rose from 62% in 2013 to 69% in 2017 (OR=1.2; p=0.095) (Fig 4). During the study period, 124,555 rabies vaccines were provided for the estimated 2,240,000 pets in Uganda, resulting in a maximum of 5.6% of pets vaccinated. Of the 6,576 monthly veterinary reports expected from the districts, only 2,517 (38%) overall were received during the study period. The percentage of expected reports received decreased from 73% in 2013 to 19% in 2017 (OR=0.67, P<0.001) (Fig 1)

**Fig 4.**
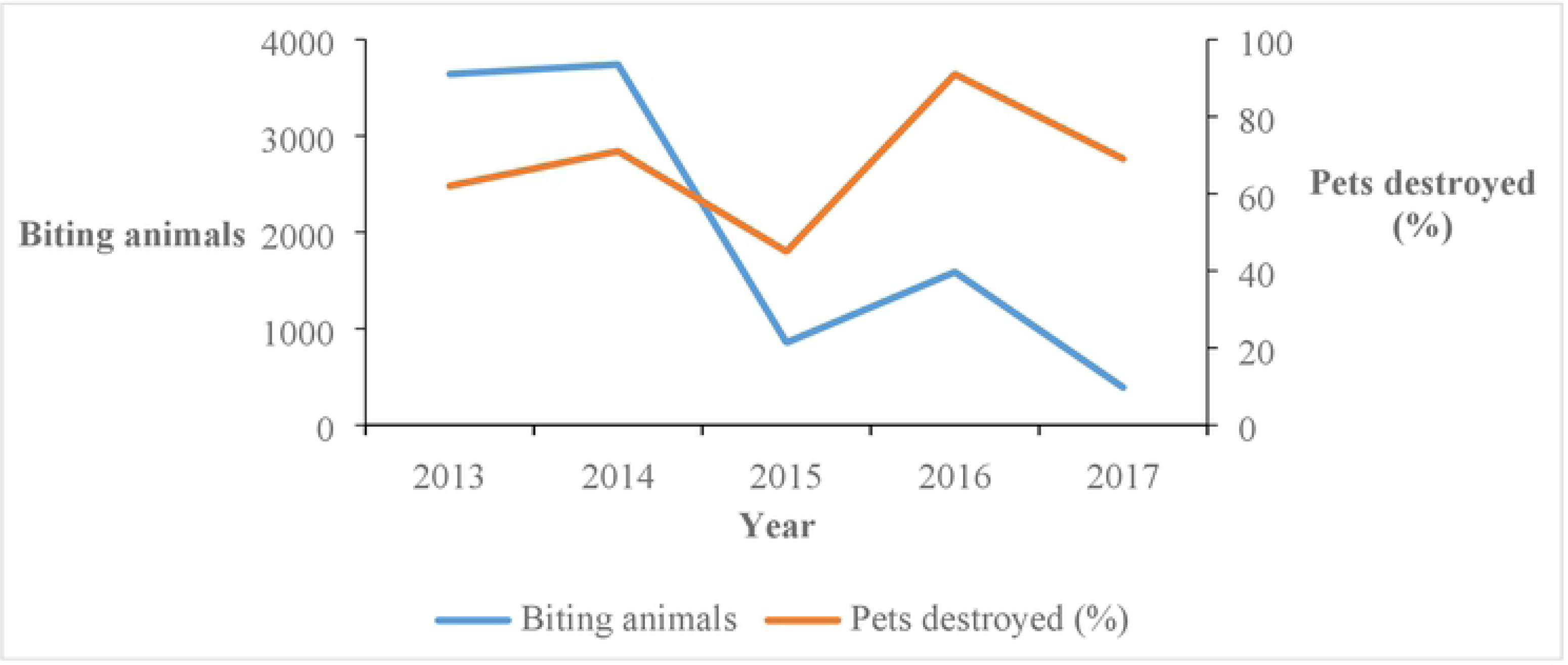
Reported trends of pets destroyed(%) among biting animals in Uganda: 2013-2017.

## Discussion

We describe veterinary surveillance data for suspected rabies in Uganda, which show an overall decline in animal bites and bite-associated human mortality in Uganda during 2013-2017. In addition, the proportion of bitten persons receiving rPEP decreased over time. Most of the animal samples tested did not test positive for rabies.

Although the data in this report appear promising for rabies reduction in Uganda, reporting rates over the study period decreased greatly. During 2013 and 2014 in Uganda, active animal disease surveillance was being supported by donors, including the Pan African Control of Epizootics (PACE) and others (18). This enabled district veterinary officers to physically bring hard-copy monthly reports to CAH, and hence to NADDEC, which improved the completeness of reporting and likely provided a more accurate picture of each of the metrics included in NADDEC rabies reporting. This support ended in 2015, which drastically affected not only rabies surveillance and reporting, but also other district-level animal disease reporting in Uganda. Other studies have also shown that passive surveillance for rabies results in underreporting (19), particularly in developing countries (20,21). Analysis of ESD rabies surveillance data from 2001-2015 found that animal bites were increasing in Uganda (14), and it is reasonable to assume that complete reporting in NADDEC might have shown a similar trend. A full evaluation of both rabies surveillance systems and the challenges to complete reporting is important to enable the system to contribute to its fullest extent in Uganda.

Regardless, findings from this study provide important information for Uganda. Vaccinating pets is one of the most important ways to prevent rabies infections among both animals and humans (7,22). However, there are clear gaps in pet vaccination in Uganda; fewer than one in ten pets were vaccinated in our analysis. This may be due in part to the limited quantities of anti-rabies vaccine procured by MAAIF for the districts (11). Both accessibility of dogs for vaccination and insufficient knowledge of dog population sizes have been shown to limit canine rabies control (23). However, rabies control can be achieved with sufficient vaccination rates: a study modelling the minimum dog vaccination coverage required for interruption of transmission of rabies in humans in Ndjamena, Chad showed that only 71% of dogs would require vaccination (24). Mass oral rabies vaccination for free-ranging dogs (OVD), which has been researched and promoted by WHO since 1988, should also be a part of the rabies control strategy (25).

While the total numbers of biting animals reported declined, the proportion of biting animals destroyed appeared to increase slightly throughout the study period. However, not all biting pets were destroyed. The accuracy of these data is unclear. Some biting pets might have escaped before the community had an opportunity to kill them; this is a common occurrence (26,27). The community might also have destroyed the pet but not reported it to the DVO, as has been reported to occur elsewhere (12). A follow up evaluation of knowledge, attitudes and practices around animal bites in the communities may be necessary to understand this issue more fully.

Most persons bitten by pets in our study received rPEP. In Uganda, rPEP is provided by government and can easily be found at high-level health centers and hospitals. However, the proportion of persons receiving rPEP in Uganda after a bite declined over the study period. The reasons for this are unclear but may relate to changes in availability at government health centers. In some areas, although rPEP is supposed to be provided free of charge, patients may be charged regardless, which can impact uptake. At least one study has shown that the costs of rPEP are typically underestimated (12). Alternately, changes in healthcare-seeking behaviors may have occurred that resulted in the decline; however, further qualitative investigation would be needed to identify these.

Furthermore, our findings indicated that approximately 1/3 of animals tested had rabies. The heads that were sent for testing likely reflect animals that had a higher pre-test probability of having rabies, thus this proportion is likely to be an overestimate of overall rabies infection among animals biting humans. In addition, among persons who did not receive rPEP, few died, suggesting that most of the animals biting might not have rabies. This is consistent with other studies that have shown that most biting animals do not have rabies (28). However, it is also possible that some animal bite victims accessed rPEP from private health facilities and were not captured in the rabies surveillance system in the veterinary sector. It is also possible that deaths due to suspected rabies could have been missed in surveillance, as rabies has a varied incubation period (three weeks to several years) and may not have been properly recognized and reported (6,10). A more detailed investigation and follow-up of persons bitten by animals, as well as active surveillance to track rPEP, could inform this issue.

Although animal bites were reported throughout Uganda, Western region had the highest incidence of animal bites during the study period. This may be due to its proximity to the Democratic Republic of Congo (DRC), which reported alarming numbers of people bitten by rabid dogs in 2013 (29,30). The thick forests of eastern DRC that border Uganda may shelter reservoirs of rabies such as jackals, and serve as sources of cross-border infections(30). In rabies elimination efforts in Uganda, this area may need special attention.

### Limitations

We were unable to link individual rPEP results to outcomes, preventing us from knowing if persons who died did or did not receive rPEP. We also did not have data about their causes of death, making it uncertain if they died from rabies. Beyond this, we did not have recent data about the number of pets in Uganda, nor about changes over time, and could only access a single value for pet numbers from the year 2008. The increase in human population in Uganda almost certainly indicates an increase in pets as well, which would have led to overestimations of vaccination rates. In addition, there was substantial underreporting, which almost certainly led to an underestimation of the magnitude of the animal bites and hence rabies in our study.

### Conclusions and recommendations

Animal bites decreased in Uganda, with western Uganda having the highest bite rate. Rabies PEP receipt among bite victims decreased over time, and overall, very few pets received anti-rabies vaccine nationwide. There was a decline in the reporting rate during 2013-2017. Evaluation of barriers to complete reporting may facilitate interventions to enhance surveillance quality. We recommended improved vaccination of pets against rabies through legislation, immediate administration of exposed humans to post-exposure anti-rabies vaccine, and sensitization of the public about the consequences of animal bites and need for urgent health care.

## List of Abbreviations

MAAIF: Ministry of Agriculture, Animal Industry and Fisheries
MoH: Ministry of Health
NADDEC: National Animal Disease Diagnostic Epidemiology Centre
OR: Odds Ratio
rPEP: Rabies Post Exposure Prophylaxis
P: probability value
PHFP: Public Health Fellowship Program

## DECLARATIONS

### Ethical approval

We got permission to conduct this investigation from the NADDEC, of the Ministry of Agriculture, Animal Industry and Fisheries. Additionally, CDC determined that this investigation was a public health emergency activity whose primary intent was to control rabies at the source (animals) such that exposure and transmission to humans is curtailed, and therefore it was classified as not research.

### Availability of data and materials

The datasets upon which our findings are based belong to NADDEC and the Uganda Public Health Fellowship Program. For confidentiality reasons the datasets are not publically available. However, the data sets can be availed upon reasonable request from the corresponding author and with permission from the NADDEC and the Uganda Public Health Fellowship Program.

### Funding and Disclaimer

This project was supported by the President’s Emergency Plan for AIDS Relief (PEPFAR) through the US Centers for Disease Control and Prevention Cooperative Agreement number GH001353–01 through Makerere University School of Public Health to the Uganda Public Health Fellowship Program, MoH. Its contents are solely the responsibility of the authors and do not necessarily represent the official views of the US Centers for Disease Control and Prevention, the Department of Health and Human Services, Makerere University School of Public Health, or the MoH. The staff of the funding body provided technical guidance in the design of the study, ethical clearance and collection, analysis, and interpretation of data and in writing the manuscript.

### Authors’ contributions

FM designed the study, and took lead in data collection, analysis, interpretation and drafting of the manuscript. DBN provided expertise on the design of the study, data collection tools, DK supervised data collection, analysis, and interpretation of the study. ARA participated in the design, analysis, and interpretation of the data; LB participated in the design, analysis, and interpretation of the data and supervised the design and drafting of the manuscript. All the authors reviewed the manuscript to ensure scientific integrity and intellectual content.

## Acknowledgements

The authors are indebted to the Uganda’s National Animal Disease Diagnostic Epidemiology Centre (NADDEC) of the Ministry of Agriculture, Animal Industry and Fisheries (MAAIF), for providing access to the rabies surveillance data and laboratory results of animal samples. We also appreciate Uganda Public Health Fellowship for the technical support during the design, analysis, and interpretation of this study. We thank PHFP cohort 2018 Fellows for the technical support during the execution of this study.

## References

1. Afakye K, Kenu E, Nyarko KM, Johnson SAM, Wongnaah F, Bonsu GK. Household exposure and animal-bite surveillance following human rabies detection in Southern Ghana. Pan Afr Med J. 2016;25(Suppl 1):12.

2. Yousaf MZ, Qasim M, Zia S, Rehman Khan M ur, Ashfaq UA, Khan S. Rabies molecular virology, diagnosis, prevention and treatment. Virology Journal. 2012 Feb 21;1(1):50.

3. Hampson K, Coudeville L, Lembo T, Sambo M, Kieffer A, Attlan M, et al. Estimating the Global Burden of Endemic Canine Rabies. PLOS Neglected Tropical Diseases. 2015 Apr 16;1(4):e0003709.

4. Tenzin T, Namgyal J, Letho S. Community-based survey during rabies outbreaks in Rangjung town, Trashigang, eastern Bhutan, 2016. BMC Infectious Diseases. 2017 Apr 17;1(1):281.

5. Sudarshan MK, Madhusudana SN, Mahendra BJ, Rao NSN, Ashwath Narayana DH, Abdul Rahman S, et al. Assessing the burden of human rabies in India: results of a national multi-center epidemiological survey. International Journal of Infectious Diseases. 2007 Jan 1;1(1):29–35.

6. Crowcroft NS, Thampi N. The prevention and management of rabies. BMJ. 2015 Jan 14;1:g7827.

7. Fooks AR, Banyard AC, Horton DL, Johnson N, McElhinney LM, Jackson AC. Current status of rabies and prospects for elimination. The Lancet. 2014 Oct 11;1(9951):1389–99.

8. Nardo PD, Gentilotti E, Vairo F, Nguhuni B, Chaula Z, Nicastri E, et al. A retrospective evaluation of bites at risk of rabies transmission across 7 years: The need to improve surveillance and reporting systems for rabies elimination. PLOS ONE. 2018 Jul 2;1(7):e0197996.

9. Shite A, Guadu T, Admassu B. Challenges of Rabies. In 2015.

10. Hooper DC. Rabies Virus. Manual of Molecular and Clinical Laboratory Immunology, Eighth Edition. 2016 Jan 1;665–73.

11. POLICY-STATEMENT-FOR-THE-MINISTRY-OF-AGRICULTURE-ANIMAL-INDUSTRY-AND-FISHERIES-FOR-THE-FINANCIAL-YEAR-2016-17.pdf [Internet]. [cited 2019 Apr 16]. Available from: http://csbag.org/wp-content/uploads/2016/05/POLICY-STATEMENT-FOR-THE-MINISTRY-OF-AGRICULTURE-ANIMAL-INDUSTRY-AND-FISHERIES-FOR-THE-FINANCIAL-YEAR-2016-17.pdf

12. The Burden of Rabies in Tanzania and Its Impact on Local Communities [Internet]. [cited 2019 Sep 5]. Available from: https://journals.plos.org/plosntds/article?id=10.1371/journal.pntd.0002510

13. Ramos J, Melendez N, Reyes F, Gudiso G, Biru D, Fano G, et al. Epidemiology of animal bites and other potential rabies exposures and anti-rabies vaccine utilization in a rural area in Southern Ethiopia. Annals of Agricultural and Environmental Medicine. 2015 Feb 24;1(1):76–9.

14. Masiira B, Makumbi I, Matovu JKB, Ario AR, Nabukenya I, Kihembo C, et al. Long term trends and spatial distribution of animal bite injuries and deaths due to human rabies infection in Uganda, 2001-2015. PLoS ONE. 2018;1(8):e0198568.

15. Kaneene J, Kabasa JD, Muleme M, Nguna J, Mafigiri R, Birungi D. Capacity building in Integrated Management of Trans-boundary Animal Diseases and Zoonoses. :12.

16. CDC - Diagnosis: Direct Fluorescent Antibody Test - Rabies [Internet]. 2019 [cited 2019 Aug 1]. Available from: https://www.cdc.gov/rabies/diagnosis/direct_fluorescent_antibody.html

17. 05_2019THE_NATIONAL_LIVESTOCK_CENSUS_REPORT_2008.pdf [Internet]. [cited 2019 Sep 14]. Available from: https://www.ubos.org/wp-content/uploads/publications/05_2019THE_NATIONAL_LIVESTOCK_CENSUS_REPORT_2008.pdf

18. PACE_booklet.pdf [Internet]. [cited 2019 Aug 23]. Available from: http://www.fao.org/ag/againfo/programmes/documents/grep/PACE_booklet.pdf

19. Taylor LH, Hampson K, Fahrion A, Abela-Ridder B, Nel LH. Difficulties in estimating the human burden of canine rabies. Acta Tropica. 2017 Jan 1;1:133–40.

20. Hampson K, Coudeville L, Lembo T, Sambo M, Kieffer A, Attlan M, et al. Estimating the Global Burden of Endemic Canine Rabies. PLOS Neglected Tropical Diseases. 2015 Apr 16;1(4):e0003709.

21. Fèvre EM, Kaboyo RW, Persson V, Edelsten M, Coleman PG, Cleaveland S. The epidemiology of animal bite injuries in Uganda and projections of the burden of rabies. Tropical Medicine & International Health. 2005 Aug 1;1(8):790–8.

22. Wallace RM, Undurraga EA, Blanton JD, Cleaton J, Franka R. Elimination of Dog-Mediated Human Rabies Deaths by 2030: Needs Assessment and Alternatives for Progress Based on Dog Vaccination. Front Vet Sci [Internet]. 2017 [cited 2019 Aug 1];4. Available from: https://www.frontiersin.org/articles/10.3389/fvets.2017.00009/full

23. Lembo T, Hampson K, Kaare MT, Ernest E, Knobel D, Kazwala RR, et al. The Feasibility of Canine Rabies Elimination in Africa: Dispelling Doubts with Data. PLOS Neglected Tropical Diseases. 2010 Feb 23;1(2):e626.

24. Zinsstag J, Lechenne M, Laager M, Mindekem R, Naïssengar S, Oussiguéré A, et al. Vaccination of dogs in an African city interrupts rabies transmission and reduces human exposure. Science Translational Medicine. 2017 Dec 20;1(421):eaaf6984.

25. Cliquet F, Guiot A-L, Aubert M, Robardet E, Rupprecht CE, Meslin F-X. Oral vaccination of dogs: a well-studied and undervalued tool for achieving human and dog rabies elimination. Veterinary Research. 2018 Jul 13;1(1):61.

26. Fekadu M. RABIES IN ETHIOPIA. Am J Epidemiol. 1982 Feb 1;1(2):266–73.

27. Gorman J. Rabies Kills Tens of Thousands Yearly. Vaccinating Dogs Could Stop It. The New York Times [Internet]. 2019 Jul 22 [cited 2019 Sep 5]; Available from: https://www.nytimes.com/2019/07/22/science/rabies-dogs-india.html

28. Medley AM, Millien MF, Blanton JD, Ma X, Augustin P, Crowdis K, et al. Retrospective Cohort Study to Assess the Risk of Rabies in Biting Dogs, 2013–2015, Republic of Haiti. Trop Med Infect Dis [Internet]. 2017 Jun 12 [cited 2019 Sep 3];2(2). Available from: https://www.ncbi.nlm.nih.gov/pmc/articles/PMC6082081/

29. Family infected after child bite in rabies horror - Democratic Republic of the Congo [Internet]. ReliefWeb. [cited 2019 Aug 23]. Available from: https://reliefweb.int/report/democratic-republic-congo/family-infected-after-child-bite-rabies-horror

30. Democratic Republic of Congo: MSF starts emergency rabies intervention [Internet]. Médecins Sans Frontières (MSF) International. [cited 2019 Aug 23]. Available from: https://www.msf.org/democratic-republic-congo-msf-starts-emergency-rabies-intervention

